# Familiar neighbours, but not relatives, enhance fitness in a territorial mammal

**DOI:** 10.1101/589002

**Authors:** Erin R. Siracusa, Stan Boutin, Ben Dantzer, Jeffrey E. Lane, David W. Coltman, Andrew G. McAdam

**Affiliations:** Department of Integrative Biology, University of Guelph, 50 Stone Rd E, Guelph, ON, N1G 2W1, CA; Centre for Research in Animal Behaviour, School of Psychology, University of Exeter, Stocker Rd, Exeter, EX4 4PY, UK; Department of Biological Sciences, University of Alberta, 116 St & 85 Ave, Edmonton, AB, T6G 2R3, CA; Department of Psychology, University of Michigan, 500 S State St, Ann Arbor, MI, 48109, USA; Department of Ecology & Evolutionary Biology, University of Michigan, 500 S State St, Ann Arbor, MI, 48109, USA; Department of Biology, University of Saskatchewan, Science Pl, Saskatoon, SK, S7N 5C8, CA; Department of Ecology and Evolutionary Biology, University of Colorado, 1900 Pleasant Street, Boulder, CO, 80309, USA

**Keywords:** dear-enemy, familiarity, fitness, kin selection, mutualism, reciprocity, red squirrel, reproductive success, senescence, social behaviour, survival, territoriality

## Abstract

**Summary:** One of the outstanding questions in evolutionary biology is the extent to which mutually beneficial interactions and kin-selection can facilitate the evolution of cooperation by mitigating conflict between interacting organisms. The indirect fitness benefits gained from associating with kin are an important pathway to conflict resolution [1], but conflict can also be resolved if individuals gain direct benefits from cooperating with one another (e.g. mutualism or reciprocity) [2]. Owing to the kin-structured nature of many animal societies, it has been difficult for previous research to assess the relative importance of these mechanisms [3–5]. However, one area that might allow for the relative roles of kin-selection and mutualistic benefits to be disentangled is in the resolution of conflict over territorial space [6]. While much research has focused on group-living species, the question of how cooperation can first be favoured in solitary, territorial species remains a key question. Using 22 years of data from a population of North American red squirrels, we assessed how kinship and familiarity with neighbours affected fitness in a territorial mammal. While living near kin did not enhance fitness, familiarity with neighbours increased survival and annual reproductive success. These fitness benefits were strong enough to compensate for the effects of aging later in life, with potential consequences for the evolution of senescence. We suggest that such substantial fitness benefits provide the opportunity for the evolution of cooperation between adversarial neighbours, offering insight into the role that mutually beneficial behaviours might play in facilitating and stabilizing social systems.

**Graphical Abstract:** 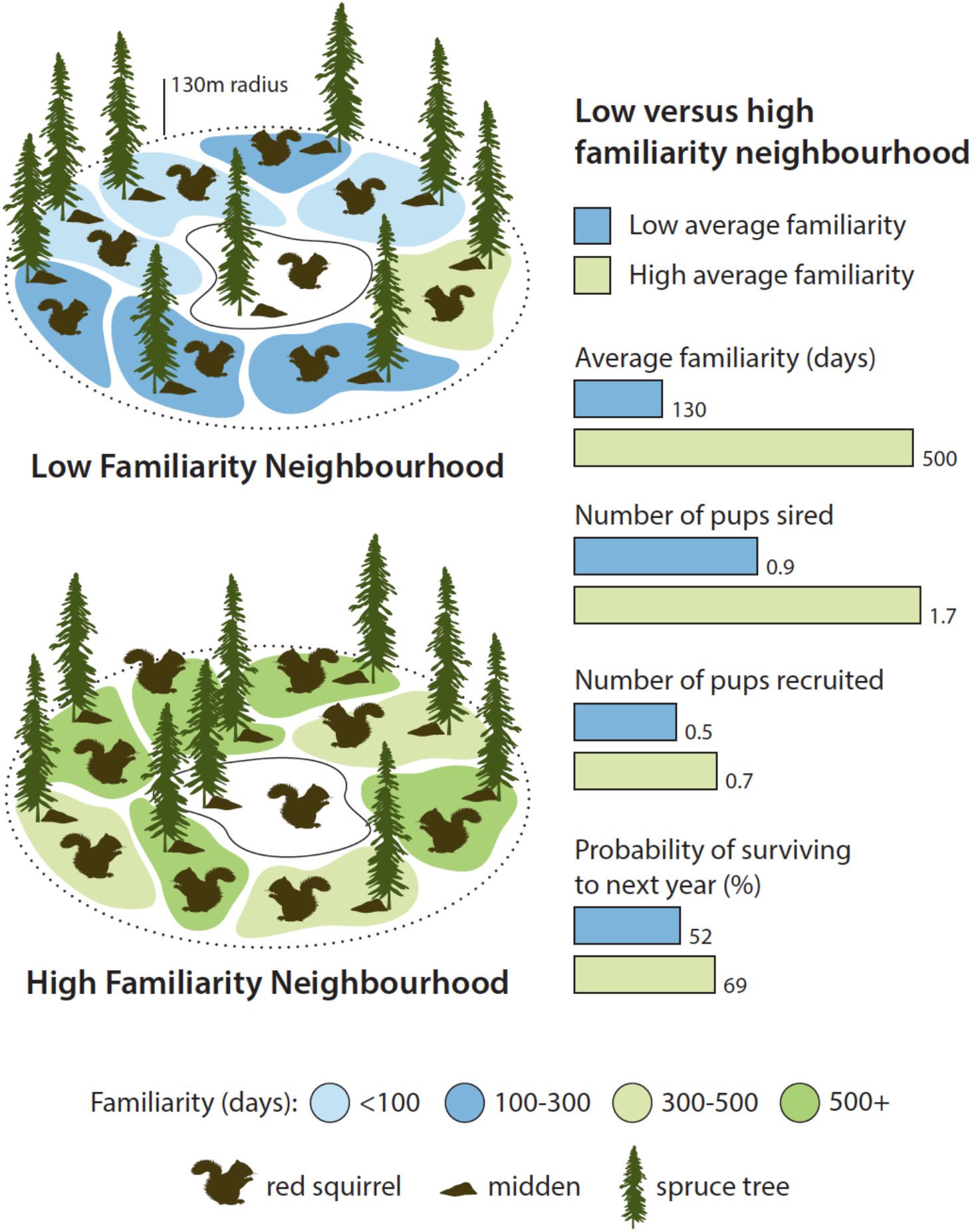

## Results and Discussion

Stable social relationships among group-living species are known to play an important role in mediating intragroup conflict and enhancing fitness [3,7,8]. For territorial species, maintaining territorial boundaries can be a costly endeavor, and while kin can help to mitigate territorial aggression and enhance fitness [9], there is also potential for stable interactions between neighbours to play an important role in helping to resolve conflict over territorial space [6,10]. It is well understood that familiarity among long-term neighbours allows for the formation of stable social relationships that can facilitate reduced aggression and territory defence, a phenomenon known as the ‘dear enemy’ effect [11,12]. Although it is not clear whether dear-enemy relationships are stabilized by reciprocity or mutualism, by minimizing negotiation of territory boundaries and alleviating costs of territoriality for both individuals, social familiarity provides a pathway to conflict resolution through mutual direct benefits.

To assess the relative importance of kinship and familiarity for resolving conflict over territorial space and thereby enhancing reproductive success and survival, we used 22 years of data from a natural population of North American red squirrels (*Tamiasciurus hudsonicus*). Red squirrels are arboreal rodents in which both sexes defend exclusive territories year-round [13]. Territories are important for overwinter survival [14] and are defended through vocalizations called ‘rattles’ [15], which are uniquely identifiable [16]. Juveniles typically disperse 100 m on average and females sometimes bequeath all or part of their territory to offspring [17]. As a result, neighbours are sometimes closely related and have been shown to exhibit affiliative behaviour with kin [18,19]. Red squirrels also have potential to establish long-term familiarity with neighbours since individuals rarely relocate to a vacant territory following natal dispersal [20]. Familiarity with neighbours is demonstrated to have important benefits for red squirrels including reduced risk of territory intrusion [21] and reduced time spent on territory defence [22].

In this study we tested whether red squirrels that were more familiar with neighbouring conspecifics had increased annual reproductive success (ARS) and a higher probability of surviving to the following year, than squirrels that were less familiar with their social neighbourhood (i.e. squirrels within a 130 m radius). Through genetic analysis and monitoring of red squirrel pups in the natal nest we were able to establish a long-term pedigree, allowing us to assess effects of social familiarity while simultaneously accounting for relatedness. We used average neighbourhood familiarity and relatedness as our metrics of interest, as previous research has demonstrated that these metrics have important effects on territory dynamics and behaviour in red squirrels [21,22]. Given that stable social connections have been demonstrated to reduce mortality risk and enhance longevity in other species [23], we also assessed whether maintaining familiar social relationships into later life might help to buffer squirrels against age-related declines in fitness. To do this, we looked for effects of familiarity on survival and reproductive success in red squirrels specifically during the ‘senescent’ period (squirrels ≥ 4 years old) [24,25].

### Effects of familiarity

Over the 22-year period (1994-2015) in which we analyzed survival and ARS, red squirrels, on average, had 13 neighbours. Average neighbourhood familiarity ranged between 0 and 1186 days (mean = 191 ± 4 days; CV = 1.04) and average relatedness ranged between 0 and 0.5 (mean = 0.05 ± 0.001; CV = 1.09). Familiarity was not strongly correlated with either relatedness (r = −0.09) or grid-wide density (r = −0.17). We found no effect of average relatedness of the social neighbourhood on male or female survival, number of pups sired annually, or number of pups recruited annually in either the full models or senescent models (all |*β*| < 0.14, all |z| < 1.48, all P > 0.13; Table 1). In contrast, familiarity with neighbouring individuals was associated with an increase in male and female survival (*β* = 0.18 ± 0.06, *z* = 2.79, *P* = 0.005; Table 1) and an increase in the annual number of pups sired (*β* = 0.18 ± 0.09, *z* = 2.04, *P* =0.04; Table 1), although there was no overall effect of social familiarity on the annual number of pups recruited by females (*β* = 0.06 ± 0.05, *z* = 1.38, *P* =0.17; Table 1). Familiarity is, however, strongly correlated with age (and therefore survival) in early life (r = 0.60), making it difficult to disentangle whether familiarity drives survival or survival drives familiarity. However, for senescent squirrels (≥ 4 years), age and familiarity are disassociated (r = −0.04; Figure S1), such that familiarity is not driven by an individual’s own survival but by the survival of its neighbours, allowing us to better disentangle these effects. When analyzing effects of familiarity in the senescent period (≥ 4 years old), we found that the benefits of social familiarity for all fitness measures were even more substantial, and were consistently at least three times greater than the effects of relatedness in the same models (Table 1). For squirrels aged 4 and older, living near familiar neighbours was associated with an increase in the probability of annual survival for both sexes (*β* = 0.45 ± 0.11, *z* = 3.40, *P* <0.001; Figure 1). For example, senescent squirrels with high familiarity with their neighbours (500 days; third quartile) had a 69% chance of surviving to the next year, while squirrels with low familiarity with their neighbours (130 days; first quartile) had a 52% chance of surviving to the next year. In the senescent period, living near familiar neighbours was associated with a substantial increase in the annual number of pups sired (*β* = 0.39 ± 0.15, *z* = 2.67, *P* = 0.008; Figure 1). Males with high familiarity with their neighbours (500 days, third quartile) sired 1.7 pups annually while males with low familiarity with their neighbours (130 days, first quartile) sired 0.9 pups annually. For females aged 4 and older, familiarity with neighbours was also associated with an increase in the annual number of pups recruited (*β* = 0.23 ± 0.09, *z* = 2.52, *P* =0.01; Figure 1). Senescent females with high familiarity with their neighbours (500 days; third quartile) recruited 0.7 pups annually compared to females with low average familiarity with their neighbours (130 days; first quartile) that recruited 0.5 pups annually.

**Table 1.**
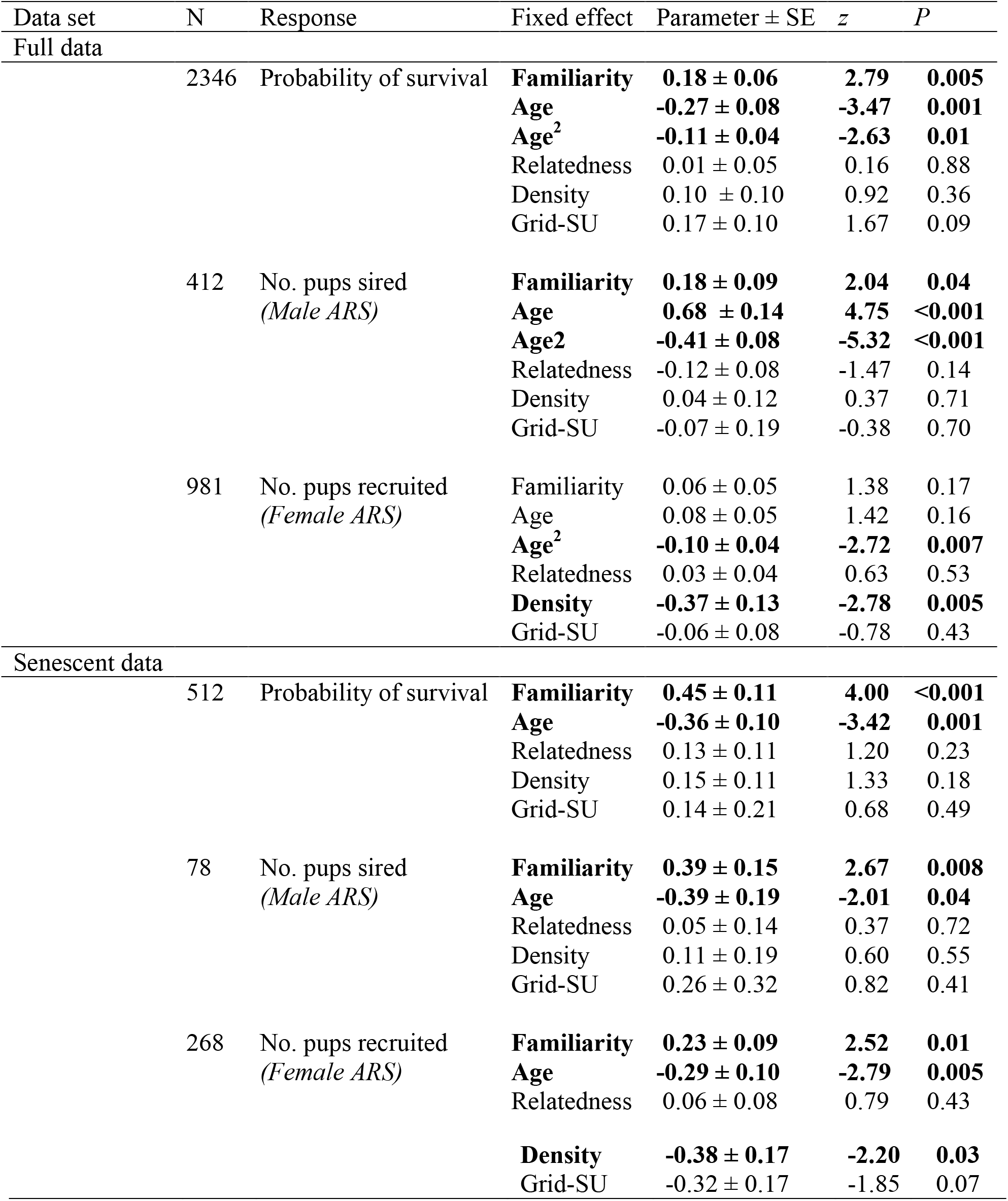
Fixed effects from annual survival, male annual reproductive success (ARS), and female ARS generalized linear mixed-effects models. Models are based on the full and senescent (≥ 4 years old) datasets are shown with significant effects indicated in bold. Regression coefficients are standardized. See also Table S1 for random effects.

**Figure 1.**
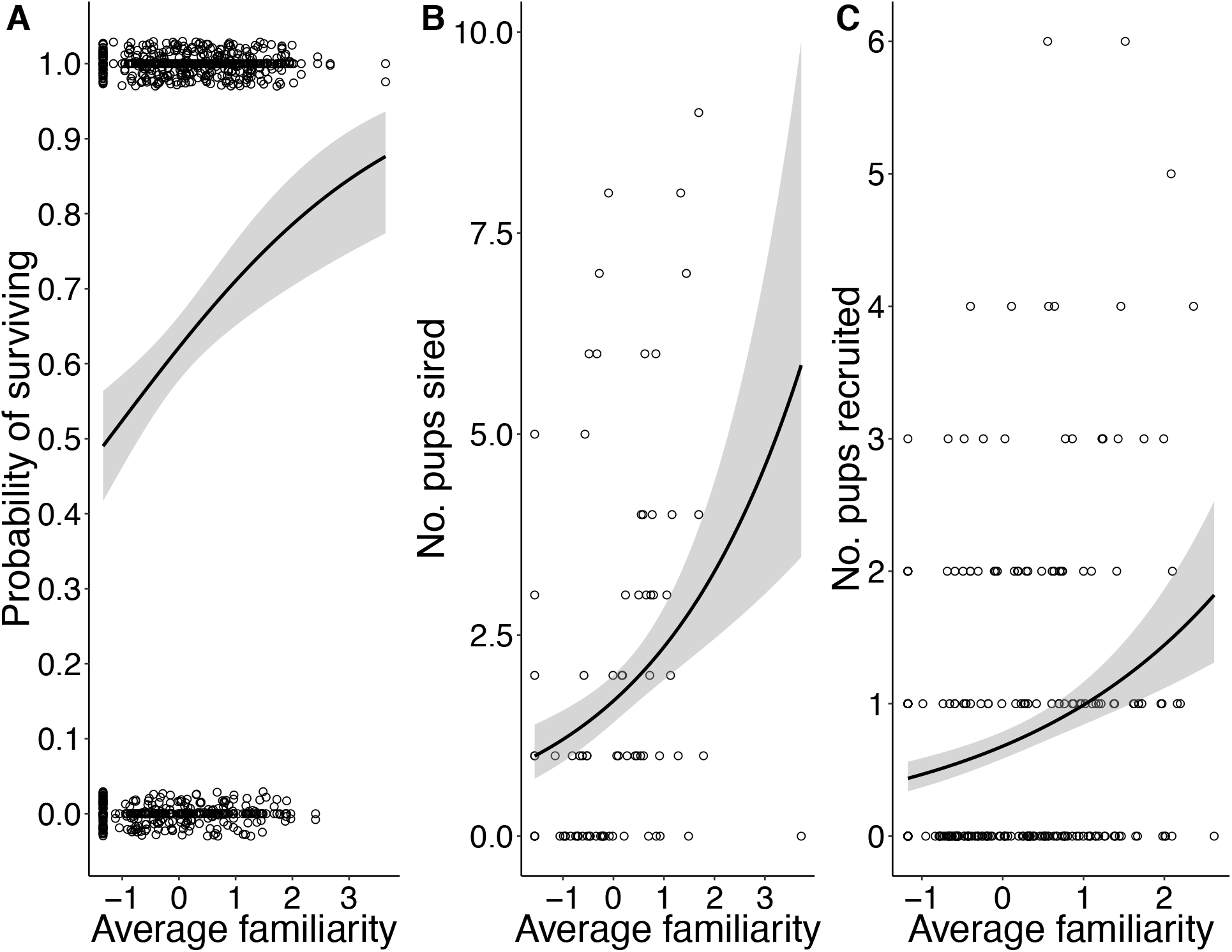
Effects of average neighbourhood familiarity on A) annual survival (N = 512), B) male annual reproductive success (ARS; N = 78), and C) female ARS (N = 268) during the senescent period (≥ 4 years old). Shaded grey bars indicate 95% confidence intervals. Values on x-axis are standardized measures of average familiarity. Points indicate raw data with a small amount of jitter introduced to show overlapping points.

An important consideration is whether variation in individual quality or habitat quality might lead to a spurious correlation between familiarity and survival when the causation is actually reversed. Familiarity, however, is not a trait of an individual but a trait of an interaction *between* individuals. Therefore, an individual with ‘good genes’ that survives well cannot obtain high familiarity without also having neighbours that survive well. For this to be possible there would need to be spatial clustering of ‘high quality’ individuals or spatial variation in resource availability or predation risk that leads to local areas of consistently high survival (and therefore high familiarity). Furthermore, in order for local areas of high habitat or individual quality to confound the familiarity-fitness relationship, spatial autocorrelation in these fitness measures must persist across multiple years (because familiarity is a product of past survival). To assess this possibility, we calculated Moran’s I for each study grid and year to test whether similarity in ARS or probability of survival was related to the spatial proximity of any two individuals. Evidence of spatial autocorrelation in any of these measures was both rare and inconsistent (Table S4). We found only one instance of spatial correlation in consecutive years (Table S4), suggesting that a spurious correlation between ARS or survival and familiarity is unlikely. When we removed the year/grid combinations with positive spatial autocorrelation from our data and reran the analyses our conclusions remained the same (Table S5). In addition, while survival and familiarity might be confounded in younger squirrels where familiarity increases with age, we demonstrated that familiarity still has substantial effects on fitness in older squirrels where age and familiarity are independent and variation in familiarity is driven by the survival of a squirrel’s neighbours rather than their own survival (Figure 1; Table 1). Finally, there is no mechanism by which reproductive success might affect familiarity. Therefore, there is substantial evidence to suggest that familiarity is a cause rather than a consequence of high fitness.

Although we do not have direct evidence for the mechanism by which familiarity with neighbours might lead to increased reproductive success, previous work in this study system has shown that social familiarity results in reduced risk of pilfering [21] and reduced time spent on territory defence [22]. Conceivably then, these direct fitness benefits may result from cooperative ‘agreements’ of non-intrusion among long-term neighbours, which might increase food stores or reduce time and energy allocated to territory defence. Given the particularly substantial effects of familiarity on male siring success, we sought to provide some mechanistic understanding of how familiarity contributes to fitness by exploring two hypotheses in male red squirrels. We hypothesized first that social familiarity might directly benefit males through increased mating success with familiar females in their social neighbourhood. Alternatively, we hypothesized that if familiarity enhances energetic resources through reduced pilferage [21,22] that this might allow male squirrels in familiar social neighbourhoods to travel farther to obtain mating opportunities and thereby increase their siring success [26,27]. Our analyses revealed that there was no association between familiarity and a male’s siring success within his social neighbourhood (≤ 130 m away; *β* = 0.12 ± 0.10, *t* = 1.25, *P* =0.21; Table 2). Instead, we found evidence that males traveled farther to mate (*β* = 19.93 ± 8.83, *t* = 2.26, *P* =0.03) and sired more pups (*β* = 0.39 ± 0.10, *z* = 3.79, *P* <0.001) outside their social neighbourhood (>130 m away) when more familiar with their neighbours (Figure 2; Table 2). This does not appear to be due to inbreeding avoidance (see Table 2). Together, these results suggest that stable social relationships with neighbours enhance male fitness by allowing them to travel farther and increase their siring success [27]. This might result from familiarity with neighbours increasing the energetic resources available to males through reduced territory defence [22] or reduced cache pilfering by neighbours [21]. While the mechanisms by which social familiarity might affect female recruitment remain unexplored, there are several pathways by which this might occur, including familiarity increasing food resources, which might lead to earlier parturition [28] or faster growth rates [29], known to be associated with enhanced juvenile survival. Alternatively the reduced time spent on defence in familiar neighbourhoods might allow females to spend more time in nest [22], and more attentive mothers have been found to have enhanced reproductive success [30]. Exploring the mechanisms by which social familiarity contributes to fitness is a viable area for future research.

**Table 2.**
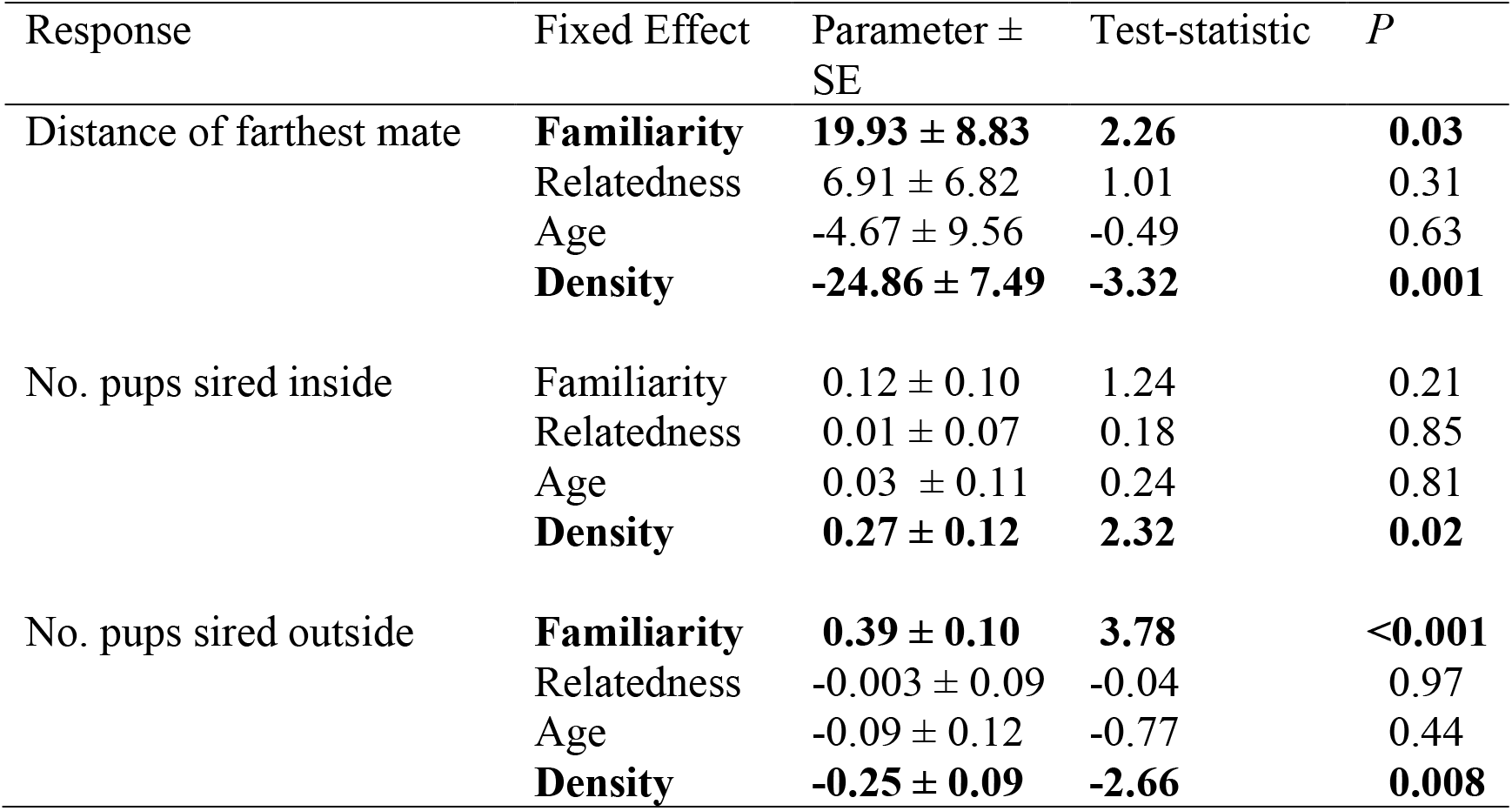
Fixed effects from models assessing effects of familiarity on (i) the farthest distance that males traveled to mate, (ii) the number of pups sired inside the neighbourhood, and (iii) the number of pups sired outside the neighbourhood (N = 199 observations over 129 males). Regression coefficients are standardized and significant effects are indicated in bold. See also Table S2 for random effects.

| Response | Fixed Effect | Parameter ± SE | Test-statistic | *P* |
| --- | --- | --- | --- | --- |
| Distance of farthest mate | **Familiarity** | **$19.93 \pm 8.83$** | **2.26** | **0.03** |
| | Relatedness | $6.91 \pm 6.82$ | 1.01 | 0.31 |
| | Age | $-4.67 \pm 9.56$ | -0.49 | 0.63 |
|  | **Density** | **$-24.86 \pm 7.49$** | **-3.32** | **0.001** |
| No. pups sired inside | Familiarity | $0.12 \pm 0.10$ | 1.24 | 0.21 |
| | Relatedness | $0.01 \pm 0.07$ | 0.18 | 0.85 |
| | Age | $0.03 \pm 0.11$ | 0.24 | 0.81 |
|  | **Density** | **$0.27 \pm 0.12$** | **2.32** | **0.02** |
| No. pups sired outside | **Familiarity** | **$0.39 \pm 0.10$** | **3.78** | **<0.001** |
| | Relatedness | $-0.003 \pm 0.09$ | -0.04 | 0.97 |
| | Age | $-0.09 \pm 0.12$ | -0.77 | 0.44 |
|  | **Density** | **$-0.25 \pm 0.09$** | **-2.66** | **0.008** |

**Figure 2.**
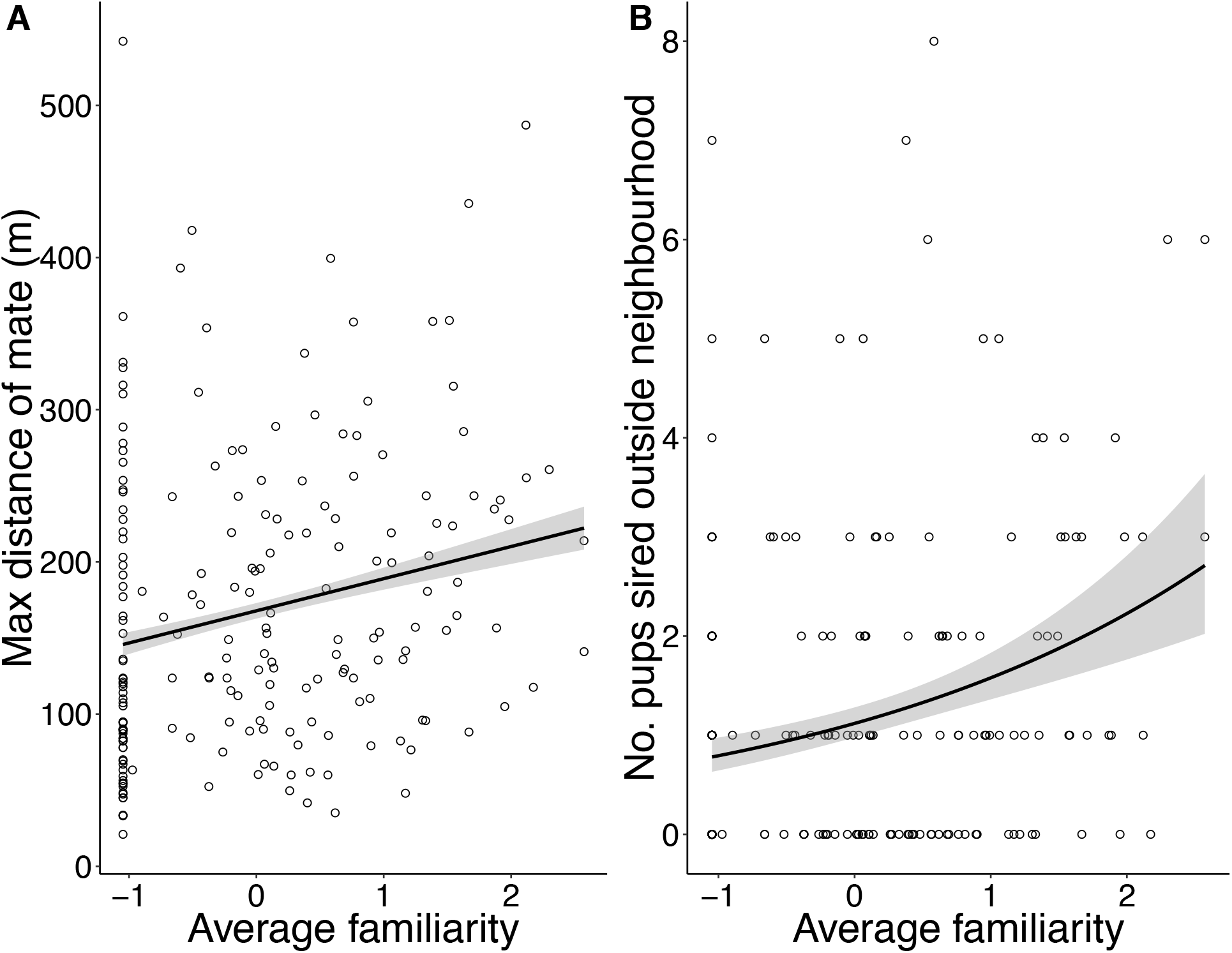
Effects of average neighbourhood familiarity on A) the distance that males traveled to mate and B) number of pups sired outside the social neighbourhood (i.e. 130 m radius; N = 199). Shaded grey bars indicate 95% confidence intervals. Values on x-axis are standardized measures of average familiarity. Points indicate raw data. See also Table 2 and S2.

### Effects on senescence

Our findings also demonstrate a previously unappreciated benefit of maintaining social relationships into later life. The magnitudes of the benefits of social familiarity in the senescent period were sufficient to offset age-related declines in survival and reproductive success. Specifically, for an average 4-year old squirrel, aging by one year decreases the probability of survival from 68% to 59%. But if that individual were to maintain all of its neighbours, such that average familiarity increased by one year (from the mean familiarity of 318 days to 683 days), the maintenance of those stable social relationships would more than compensate for the change in age, resulting in an increase in survival probability from 68% to 74%. However, despite the large individual benefits of increased social familiarity late in life, few squirrels currently enjoy this fitness advantage. For example, only 4% of 4-year olds surviving to age 5 maintain all of their neighbours, and mean familiarity does not continue to increase with age through the senescent period (Figure S1). As a result, our observed fitness increases in later life associated with social stability are not currently widespread enough to affect the decline in the force of natural selection with age and the arrival of the mutational ‘wall of death’ [31] (Figure S2).

Nevertheless, such individual fitness benefits of familiarity provide strong incentive for squirrels to reduce turnover in their social neighbourhood. While squirrels compete with territorial neighbours for resources [32], space [13], and reproductive success [33], here we have shown that individuals also benefit from the enhanced survival of neighbours. The idea that territorial neighbours might engage in cooperative behaviours to prevent neighbourhood turnover has been theorized [34]. Here our documented benefits of social familiarity provide the opportunity for the evolution of cooperative behaviour toward otherwise adversarial neighbours. Specifically, while the loss of an unfamiliar neighbour would have no effect on survival probability, the loss of a neighbour with 6 years of familiarity, although rare, would decrease the owner’s probability of survival by 7%. A squirrel should, therefore, be willing to engage in behaviours that ensure their neighbour’s survival as long as this does not reduce the actor’s survival by more than 7%. This scope for cooperation is comparable to kin selection interactions between first cousins once removed (r = 0.06) [1]. It is possible then that reduced aggression when interacting with familiar neighbours (i.e. the dear-enemy phenomenon) [12] is a mechanism that not only serves to reduce an individual’s own time and energy spent on defence, but also enhances the survival of familiar neighbours, which provides indirect fitness returns through increased social familiarity. In red squirrels, reduced territory defence toward familiar individuals [22] may convey survival benefits to neighbours through reduced aggression or lowered defense of food resources. Red squirrels have also been known to nest communally during cold temperatures [18], which might enhance neighbour survival. Other mechanisms by which red squirrels might reduce neighbourhood turnover such as predator alarm calling, or forming defensive coalitions [e.g. 34] to prevent a territory takeover, have yet to be tested. Ultimately, if cooperative behaviours enhancing the survival of familiar neighbours were to become widespread enough that mean social familiarity increased with age across the population, then this socially driven increase in fitness late in life could influence the force of selection and consequently the evolution of senescence (Figure S2).

### Effects of relatedness

Given that there is evidence of affiliative behaviour among kin in this study system [18,19], in addition to localized dispersal and bequeathal of territory space to offspring [17], we expected that relatedness among neighbours might play an important role in red squirrel survival and reproductive success. Additionally, we were interested in the possibility that relatedness with neighbours might help to mitigate the consequences of living with unfamiliar neighbours, as seen in other systems [6], However, we found no evidence to suggest that kin provide fitness benefits for red squirrels (Table 1), and there were no significant interactions between relatedness and familiarity in any of our models. It is possible that the observed low relatedness among neighbours (average neighbourhood relatedness = 0.05 ± 0.001), which likely occurs as the result of low overwinter juvenile survival [35], could have constrained the potential for kin selection to act at this scale. However, our findings do not mean that neighbour relatedness has no biologically meaningful effects on red squirrel fitness, but simply that the scale at which we measured these effects did not exhibit any influence of kinship. If, for instance, the primary benefits of kinship come from sharing resources such as food [21] or nests [18], relatedness of nearest neighbours (rather than all neighbours within 130 m) may be a more relevant measure to assess whether red squirrels obtain benefits from kin. Future work should therefore attempt to use other measures of the social environment to offer additional insight as to the relative importance of kin-selection in this system. Regardless, our results demonstrate that stable social relationships are associated with increased fitness in red squirrels, independent of kinship, suggesting that inclusive fitness benefits are not a necessary prerequisite to facilitate cooperation between social partners.

## Conclusion

Although interactions among territorial individuals are fundamentally defined by competition for food, mates, or other resources, here we have shown that stable social relationships can provide important fitness benefits that might feed back to limit these competitive interactions and maintain strong affiliative relationships among unrelated territorial neighbours. Similar fitness benefits of established relationships with neighbours have also been demonstrated in territorial birds such as red-winged blackbirds [36], song sparrows [37], and great tits [10]. Our findings, in conjunction with these previous studies, demonstrate that the benefits of stable social relationships are not exclusive to group-living organisms but can extend to solitary species, suggesting a shared evolutionary importance of stable relationships across a range of social complexities. The strength of the effects of familiarity, particularly in the senescent period, was surprising. We suggest that by increasing the force of selection later in life, stable social relationships have the potential to lead to the evolution of slower rates of aging, although these late-life social benefits are not currently widespread enough to affect senescence in red squirrels. Importantly, our results indicate that intraspecific conflict can be mitigated through mutualistic pathways, with resulting fitness benefits, independent of kinship effects [3,6]. Consequently, stable social relationships with neighbours may be an important and overlooked contributor to fitness in territorial species, and may offer insight into the relative importance of mutually beneficial behaviours for facilitating conflict resolution.

## Acknowledgments

We thank the Champagne and Aishihik First Nations for allowing us to conduct our research within their traditional territory and thank Agnes MacDonald and her family for long-term access to her trap line. We are grateful to all of the field technicians who have contributed to the long-term KRSP database over the years, and J. Kapitain at KAP Design for assistance with figure design. This research was supported by Discovery Grants and Northern Research Supplements from the Natural Sciences and Engineering Research Council of Canada (A.G.M.: RGPIN-2015-04707; S.B.: RGPIN-2014-05874; J.E.L.: RGPIN-2014-04093, RGPNS-2014-459038; D.W.C.: RGPIN-2011-312207), as well as funding from the University of Michigan (B.D.), and Grants-in-Aid of research from the American Society of Mammalogists and the Arctic Institute of North America (E.R.S.).

## Author Contributions

E.R.S. and A.G.M. conceived of and designed the study. E.R.S. conducted statistical analyses and wrote the manuscript with support from A.G.M. All authors contributed to field logistics, data collection and maintenance, and assisted with review and editing of the manuscript.

## Declaration of Interests

The authors declare no competing interests.

## STAR Methods

### Resource Availability

#### Lead Contact

Further information and requests for resources and reagents should be directed to and will be fulfilled by the Lead Contact, Erin Siracusa.

#### Materials Availability

This study did not generate new unique reagents.

#### Data and Code Availability

Data are available from the Figshare Repository (https://doi.org/10.6084/m9.figshare.8813564).

### Experimental Model and Subject Details

We studied a wild population of North American red squirrels located in the southwest Yukon, Canada (61° N, 138° W). This population has been followed since 1987 on two 40-ha study grids separated by the Alaska Highway (‘Kloo’ and ‘Sulphur’). We monitored squirrels annually from March to August and used a combination of live-trapping and behavioural observations to assess territory ownership, track reproduction and survival, and determine offspring recruitment [17,25]. We trapped squirrels using Tomahawk live-traps (Tomahawk Live Trap Co., Tomahawk, Wisconsin, USA) baited with peanut butter. During their single-day oestrous, female red squirrels mate with multiple males and produce multiply-sired litters [38], with an average of three pups per litter [39]. We monitored the reproductive statuses and parturition dates of females through palpation, mass changes, and evidence of lactation. After parturition, we used radio telemetry and/or behavioural observations to follow females to their nests. We fitted pups with unique alphanumeric metal ear tags (Monel #1; National Band and Tag, Newport, KY, USA) at 25 days old, allowing us to follow individuals throughout their lifetimes.

#### Ethical guidelines

This study required trapping individuals using Tomahawk live traps to determine territory ownership and assess reproductive status. Traps were checked every 60-90 min and squirrels were never left in a trap for longer than 120 min. We also entered the natal nest when pups were 1-2 days old and 25 days old to collect DNA, measure pup growth, and tag individuals. We returned the pups to the natal nest immediately after processing to minimize time spent away from the dam. These procedures had no detectable negative effects on the survival or welfare of the study animals. This research was approved by the University of Guelph Animal Care Committee (AUP number 1807).

### Method Details

#### Measuring familiarity

We completed a full census of the population twice annually in May and August and determined territory ownership using the aforementioned methods. Both male and female red squirrels defend exclusive territories year round, which are centered around a larder hoard of food resources called a ‘midden’ [13]. We defined each squirrel’s social neighbourhood to be all conspecifics whose middens (i.e. the center of the territory) were within a 130 m radius of the focal squirrel’s midden. One hundred and thirty meters is the farthest distance that red squirrel territorial vocalizations (‘rattles’) are known to be detectable to the human ear [40], and is similar to the distance at which red squirrels have been demonstrated to be most responsive to changes in local density (150 m) [41]. This suggests that 130 m is a reasonable measure of the distance at which squirrels can receive and respond to information about their social environment. We measured the pairwise familiarity between the territory owner and each neighbour as the number of days that both individuals had occupied their current territories within 130 m of each other. We then averaged all pairwise measures of familiarity to obtain a measure of each individual’s average familiarity with its neighbours. For all analyses we used average familiarity values calculated in May, as this aligned most closed with the reproductive season. The timing of our semi-annual censuses meant that we could only update each squirrel’s familiarity with its neighbours twice per year. Therefore, pairwise familiarities could increase from one year to the next in increments of either 273 days (new neighbour in August survived until May), or 365 days (neighbour survived from previous May until current May). Average familiarity, however, could take on many possible values because pairwise familiarities were averaged across all neighbours.

#### Measuring relatedness

We temporarily removed juveniles from their natal nest immediately following parturition to weigh, sex, and take ear tissue samples. Observing pups in the natal nest also allowed us to assign maternity with certainty. Starting in 2003, we determined paternity by genotyping all individuals at 16 microsatellite loci [42] and assigned paternity with 99% confidence using CERVUS 3.0 [43]. Genotyping error based on known mother-offspring pairs was 2%. Additional details on the genetic methods can be found in Lane et al. (2008). Using the established multigenerational pedigree, we calculated the coefficient of relatedness between the territory owner and each neighbour. We averaged all pairwise measures of relatedness to provide a measure of the average relatedness between each squirrel and all of its neighbours.

#### Fitness measures

For individuals alive between 1994 and 2015, we measured annual survival as the probability of surviving to the following year (*N* = 2346 records over 1009 individuals). Given that long-term monitoring of individuals in this population first began in 1987, we excluded data prior to 1994 to ensure accurate measurement of familiarity between neighbours. Our ability to successfully redetect adults in the population each year is estimated to be 1.0 [44].

We measured female annual reproductive success (ARS) as the number of pups recruited each year (i.e. surviving overwinter). We included data from 450 breeding females between 1994 and 2015 (*N* = 981 records). All females with a known parturition date were considered to have bred; all other non-breeding females were excluded from analysis. Most non-breeding females were yearlings that were too young to breed. Since red squirrel males do not contribute to the raising of pups and the male’s social environment is, therefore, unlikely to affect pup recruitment, we defined male ARS as the number of pups sired each year. We used data from 207 males between 2003 and 2014 (*N* = 412 records). We excluded male reproductive success outside of this timeframe because paternity data was not available prior to 2003 and after 2014. For all analyses we only used data for adults (≥ 1 year old) whose age could be assigned with certainty (i.e. individuals tagged in nest as juveniles).

It is possible that our ability to accurately assess recruitment is somewhat confounded by dispersal beyond the borders of our study grids [45]. On average, juveniles typically disperse less than 100 m [17,46], and estimates of juvenile survival do not differ between the center and edge of our study grids, which might be expected if dispersing juveniles were mistakenly considered to have died [25]. However, 37% of our population is comprised of immigrants from outside our study grids [45], which suggests that some juvenile recruitment outside the study area is likely to have been missed by our methods.

### Quantification and Statistical Analysis

#### Annual survival and ARS

To assess the effects of familiarity on ARS and survival, we used generalized linear mixed effects models (GLMMs) with a BOBYQA optimizer. We used a binomial distribution (logit-link) to assess probability of annual survival and a Poisson distribution (log-link) for male and female ARS. For all models we fitted average familiarity, average relatedness, a linear and quadratic term for age, and grid-wide density as continuous predictors and included study grid as a categorical fixed effect (Table 1). For the survival models we checked for an effect of sex, but sex was not significant and so was not considered further. We included a random effect of squirrel ID and year to account for repeated measures of individuals and temporal variation in resource availability, respectively (Table S1). To account for inherent spatial structure in our data, we grouped squirrels into 150 m squares (within each year) based on their known spatial locations and included this ‘spatial ID’ as a random effect in all of our models. Our results remained unchanged even when this spatial grouping term was defined to be a 75 m or 300 m square (see Table S3). Fitting separate spatial autocorrelation terms for each year would have been ideal, however, it was not possible to obtain convergence from our data using this method. To account for the fact that familiarity and age are strongly correlated in early, but not later, life (Figure S1), we also fit models for just the senescent period (≥ 4 years old). Models for the senescent period included the same covariates as the full models described above, but only included a linear effect of age. Finally, we conducted an additional analysis to test for spatial autocorrelation by calculating Moran’s I (ape package version 5.1) [47] for each study grid and year (Table S4). We removed the year/grid combinations with positive spatial autocorrelation from our data and reran the analyses above (Table S5).

#### Mechanisms underlying male reproductive success

We conducted a post-hoc analysis to reveal potential mechanisms by which familiarity with neighbours might lead to increased male reproductive success. We had two hypotheses: 1) that familiarity with neighbours leads to increased energetic resources, allowing males to travel farther to mate or 2) that familiarity with females provides males with more mating opportunities within the males’ social neighbourhood. To test hypothesis one, we assessed whether familiarity with neighbours affected the farthest distance that a male traveled to mate using a linear mixed effect model (LMM) with average familiarity, average relatedness, age, and grid density as continuous fixed effects and squirrel ID and year as random effects (Table 2, S2). To test hypothesis two, we measured whether average familiarity affected the number of pups that a male sired inside or outside his neighbourhood. We fitted both these models using a Poisson GLMM with the same fixed and random effects structure as above (Table 2, S2).

#### Force of selection

If the fitness benefits of stable social relationships were sufficiently widespread in the red squirrel system then an increase in the mean number of reproductive opportunities later in life associated with social familiarity would lead to a less rapid decline in the force of natural selection with age. This would delay the arrival of the ‘wall of death’, a spike in mortality caused by the accumulation of deleterious alleles, leading to the evolution of slower rates of aging [31]. We conducted a post-hoc analysis to explore this possibility in red squirrels by creating a ‘simulated’ population where social interactions were excluded from the calculation of age-specific survival and fecundity (Figure S2). To do this, we fitted the same models as above, except that we included both breeding and non-breeding females and the response variable for females was now number of pups born rather than number recruited. For the first set of models we included age, a quadratic term for age, density and grid as fixed effects, as well as squirrel ID and year as random effects. We extracted the parameter estimates for age in each of these models and used those parameters to estimate age-specific fecundity and survival for females and males separately. This represented our ‘observed’ population, and provided us with the estimates of age-specific fecundity and survival as observed and as potentially confounded by social familiarity among squirrels. We then measured what the effects of age would have been had there not be been variation in familiarity by including familiarity as a continuous predictor in the same models described above, and thus statistically controlling for its effects. Our parameter estimates for age now represented the effects of age in the absence of social effects. We again used these parameters to estimate age-specific fecundity and survival for both sexes. We calculated force of selection separately for males and females in our ‘observed’ population and ‘simulated’ population (where the benefits of social relationships were statistically excluded) by using the following formula [48]:

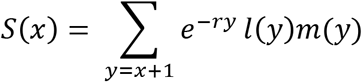

Here *S*(*x*) represents the force of selection against a deleterious allele in the population, where *r* is the intrinsic rate of increase, *l*(*y*) and *m*(*y*) are the survivorship and fecundity functions and *y* is used to sum up the net expected reproduction over all ages after age *x*.

#### Data analysis

We conducted statistical analyses using R version 3.4.1 [49] and fitted models using the lme4 package (version 1.1-13) [50]. We obtained *P* values for the LMMs using the package lmerTest (version 3.0-1) [51]. For all models, we checked for significant non-linearities between the predictor and response variables by fitting generalized additive models. For GLMMs we checked for overdispersion by assessing whether the sum of squared Pearson residuals approximated a Chi-squared distribution with *N-P* degrees of freedom [52]. Of the 1009 squirrels used in this analysis 12 individuals were not in the pedigree, meaning there were 25 neighbourhoods for which we could not calculate relatedness with neighbours. We assigned these neighbourhoods the mean average neighbourhood relatedness (r = 0.05). (Results remained the same even when these 25 neighbourhoods were removed from the analysis). To facilitate direct comparison of effect sizes [53] we standardized all continuous predictors to a mean of zero and unit variance. We estimated all effect sizes by using first and third quartile familiarity values while holding all other variables constant at their mean and using ‘Kloo’ as the study grid. We present all means ± SE, unless otherwise stated, and consider all differences statistically significant at *α* < 0.05.

## Notes

https://doi.org/10.6084/m9.figshare.8813564

### Competing Interest Statement

The authors have declared no competing interest.

### Summary of Updates

Revisions to text made to enhance clarity of the manuscript

