## Supplementary Materials for "Familiar neighbours, but not relatives, enhance fitness in a territorial mammal"

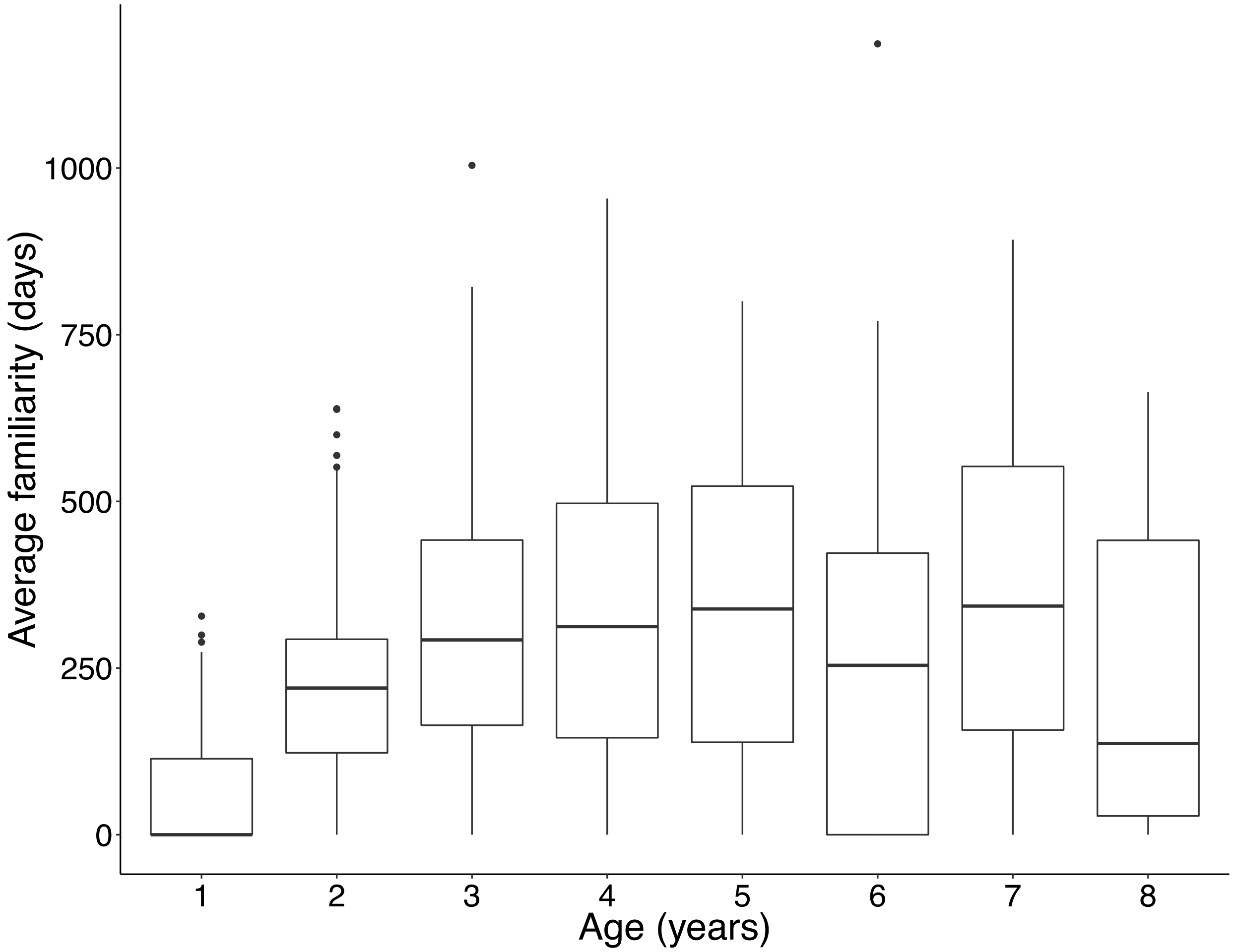


**Figure S1. Relationship between red squirrel age and the average familiarity of the social neighbourhood (conspecifics within 130 m), related to STAR Methods.** Before age four, age and familiarity are strongly correlated (Pearson r = 0.60). However, in the senescent period (≥ 4 years old) familiarity in squirrels is driven by other demographic factors in the population such that age and familiarity become decoupled (Pearson r = -0.04).

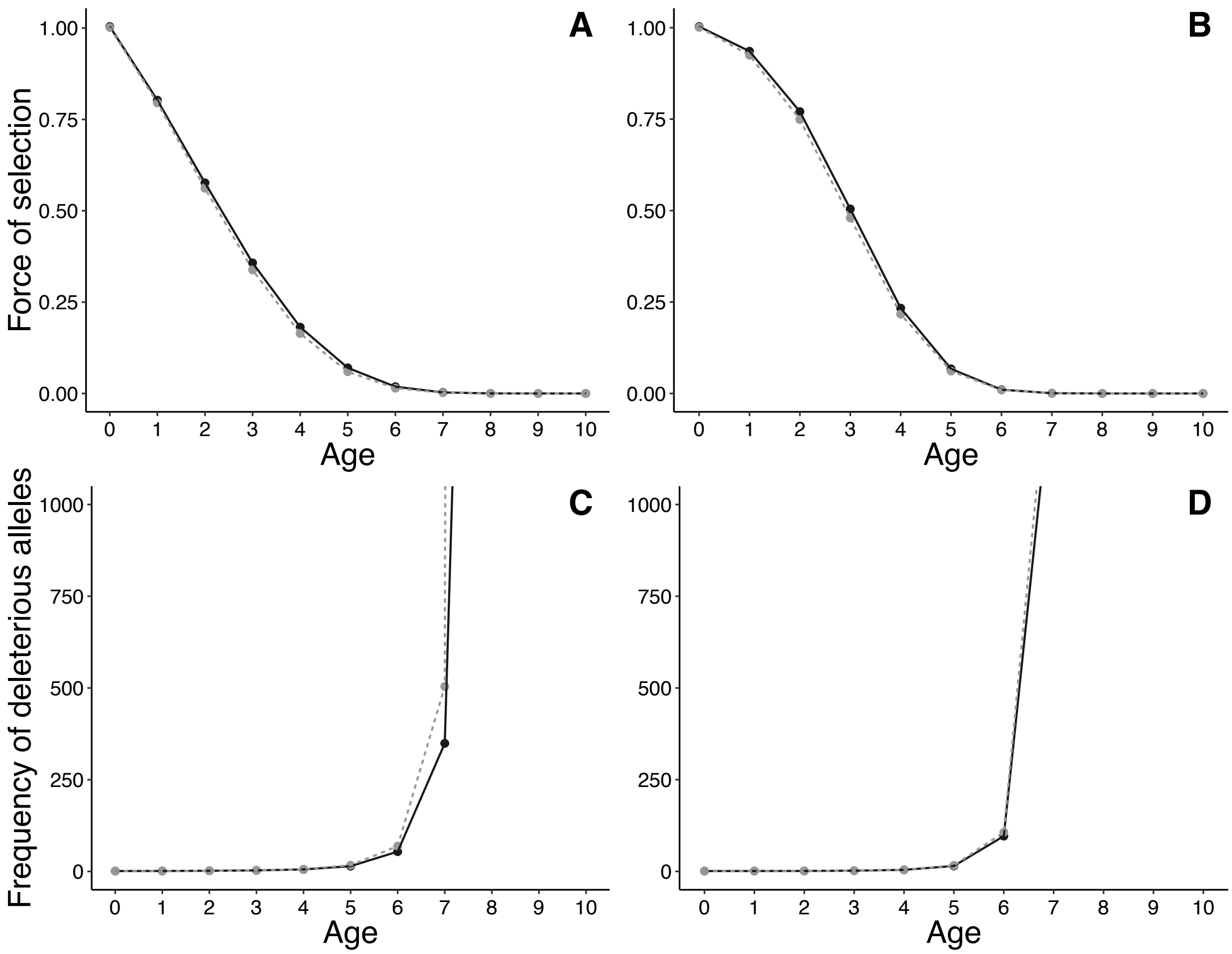


**Figure S2. Force of selection and the 'wall of death' for female and male red squirrels in our 'observed' population (solid line) and 'simulated' population (dashed grey line), related to STAR Methods.** The top two panels of this figure show Hamilton's force of selection ($S(x)$) by age for A) females and B) males. The force of selection falls to zero as the remaining survival-weighted reproduction for each sex declines. The bottom two panels of this figure show the inverse of the force of selection as an indicator of age-specific mortality for C) females and D) males. The rate of accumulation of deleterious alleles in the population is expected to be proportional to $1/S(x)$. The rapid increase in deleterious alleles observed when the force of selection drops to zero is referred to as the 'wall of death'. The solid black lines show the squirrel population under observed conditions where squirrels experience the social benefits of familiar neighbours. The dashed grey lines show the 'simulated' squirrel population without the benefits of social relationships. Under observed conditions, where red squirrels experienced the social benefits of familiar neighbours, the force of selection was marginally stronger than in a ‘simulated’ population where social interactions were excluded from the calculation of age-specific survival and fecundity (A, B). However, this ultimately had no influence on when the wall of death occurred (C, D). This is likely due to the fact that currently, in red squirrels, familiarity does not increase linearly with age (see Figure S1), meaning that while some individuals experience stable social environments and their associated fitness benefits, others have substantial turnover in neighbours and as a result have reduced survival and reproductive success. Given that not all squirrels experience the benefits of familiarity in old age, this limits the extent to which social relationships might currently affect patterns of senescence at the population level.

| Data set | Response | Random effect | Variance | $\chi^2$ | *P* |
|---|---|---|---|---|---|
| Full data | | | | | |
| | Probability of survival | Squirrel ID | <0.01 | <0.01 | >0.999 |
| | | **Year** | **0.18** | **29.00** | **<0.001** |
| | | Spatial ID | 0.07 | 0.55 | 0.46 |
| | No. pups sired *(Male ARS)* | **Squirrel ID** | **0.47** | **35.60** | **<0.001** |
| | | **Year** | **0.08** | **6.19** | **0.01** |
| | | **Spatial ID** | **0.44** | **39.28** | **<0.001** |
| | No. pups recruited *(Female ARS)* | Squirrel ID | 0.03 | 0.69 | 0.40 |
| | | **Year** | **0.34** | **128.12** | **<0.001** |
| | | Spatial ID | 0.01 | 0.08 | 0.78 |
| Senescent data | | | | | |
| | Probability of survival | Squirrel ID | <0.01 | <0.01 | >0.999 |
| | | Year | 0.06 | 0.60 | 0.44 |
| | | Spatial ID | 0.27 | 1.24 | 0.26 |
| | No. pups sired *(Male ARS)* | **Squirrel ID** | **0.45** | **5.96** | **0.01** |
| | | Year | <0.01 | <0.01 | >0.999 |
| | | **Spatial ID** | **0.37** | **4.50** | **0.03** |
| | No. pups recruited *(Female ARS)* | Squirrel ID | <0.01 | <0.01 | >0.999 |
| | | **Year** | **0.30** | **18.84** | **<0.001** |
| | | Spatial ID | 0.11 | 1.31 | 0.25 |

**Table S1. Random effects from annual survival, male annual reproductive success (ARS), and female ARS generalized linear mixed-effects models, related to Table 1.** Significance assessed using a log-likelihood ratio test (LRT) with one degree of freedom to compare models with and without the listed random effect. Significant effects are indicated in bold.

| Response | Random Effect | Variance | $\chi^2$ | *P* |
|---|---|---|---|---|
| Distance of farthest mate | Squirrel ID | 2415 | 2.92 | 0.09 |
| | Year | <0.01 | <0.01 | >0.999 |
| No. pups sired inside | **Squirrel ID** | **0.20** | **11.60** | **<0.001** |
| | **Year** | **0.08** | **6.21** | **0.01** |
| No. pups sired outside | **Squirrel ID** | **0.41** | **28.22** | **<0.001** |
| | Year | <0.01 | <0.01 | >0.999 |

**Table S2. Random effects from models assessing effects of familiarity on (i) the farthest distance that males travels to mate, (ii) the number of pups sired inside the neighbourhood, and (iii) the number of pups sired outside the neighbourhood (N = 199 observations over 129 males), related to Table 2.** Significance assessed using a log-likelihood ratio test (LRT) with one degree of freedom to compare models with and without the listed random effect. Significant effects are indicated in bold.

| Spatial term | Model | N | Fixed effect | Parameter ± SE | *z* | *P* |
|---|---|---|---|---|---|---|
| 75 m | Survival | 512 | **Familiarity** | **0.43 ± 0.11** | **3.95** | **<0.001** |
| | | | **Age** | **-0.34 ± 0.10** | **-3.41** | **<0.001** |
| | | | Relatedness | 0.11 ± 0.10 | 1.15 | 0.25 |
| | | | Density | 0.15 ± 0.11 | 1.36 | 0.17 |
| | | | Grid-SU | 0.15 ± 0.19 | 0.75 | 0.45 |
| | Male ARS | 78 | **Familiarity** | **0.39 ± 0.15** | **2.62** | **0.009** |
| | | | Age | -0.35 ± 0.19 | -1.87 | 0.06 |
| | | | Relatedness | 0.08 ± 0.14 | 0.54 | 0.59 |
| | | | Density | 0.12 ± 0.19 | 0.63 | 0.53 |
| | | | Grid-SU | 0.27 ± 0.30 | 0.92 | 0.36 |
| | Female ARS | 268 | **Familiarity** | **0.22 ± 0.09** | **2.55** | **0.01** |
| | | | **Age** | **-0.28 ± 0.10** | **-2.84** | **0.005** |
| | | | Relatedness | 0.06 ± 0.07 | 0.85 | 0.40 |
| | | | **Density** | **-0.37 ± 0.17** | **-2.17** | **0.03** |
| | | | Grid-SU | -0.32 ± 0.16 | -1.94 | 0.05 |
| 300 m | Survival | 512 | **Familiarity** | **0.43 ± 0.11** | **4.08** | **<0.001** |
| | | | **Age** | **-0.34 ± 0.10** | **-3.47** | **<0.001** |
| | | | Relatedness | 0.11 ± 0.10 | 1.16 | 0.25 |
| | | | Density | 0.15 ± 0.11 | 1.36 | 0.17 |
| | | | Grid-SU | 0.14 ± 0.20 | 0.73 | 0.46 |
| | Male ARS | 78 | **Familiarity** | **0.36 ± 0.15** | **2.50** | **0.01** |
| | | | Age | -0.32 ± 0.18 | -1.78 | 0.08 |
| | | | Relatedness | 0.12 ± 0.14 | 0.80 | 0.42 |
| | | | Density | 0.08 ± 0.19 | 0.45 | 0.65 |
| | | | Grid-SU | 0.22 ± 0.33 | 0.68 | 0.50 |
| | Female ARS | 268 | **Familiarity** | **0.23 ± 0.09** | **2.69** | **0.007** |
| | | | **Age** | **-0.28 ± 0.10** | **-2.86** | **0.004** |
| | | | Relatedness | 0.06 ± 0.08 | 0.78 | 0.44 |
| | | | **Density** | **-0.36 ± 0.17** | **-2.10** | **0.04** |
| | | | Grid-SU | -0.32 ± 0.16 | -2.00 | 0.05 |

**Table S3. Results from senescent (≥ 4 years old) models when including a 75 m x 75 m spatial grouping term or a 300 m x 300 m spatial grouping term to account for potential spatial autocorrelation in the data, related to STAR Methods.** Results are consistent with the findings presented in the manuscript where a 150 m x 150 m spatial grouping term was included in the models. Significant effects indicated in bold. Regression coefficients are standardized.

| Response Variable | Grid | Year | Observed | Expected | SD | *P* |
|---|---|---|---|---|---|---|
| No. pups recruited | | | | | | |
| | KL | 1994 | -0.08 | -0.05 | 0.04 | 0.75 |
| | | 1995 | 0.01 | -0.04 | 0.05 | 0.17 |
| | | 1996 | -0.03 | -0.03 | 0.03 | 0.51 |
| | | **1997** | **0.07** | **-0.04** | **0.04** | **0.01** |
| | | 1998 | -0.01 | -0.02 | 0.03 | 0.29 |
| | | 1999 | -0.06 | -0.03 | 0.03 | 0.82 |
| | | 2000 | -0.03 | -0.03 | 0.04 | 0.46 |
| | | **2001** | **0.05** | **-0.03** | **0.03** | **0.005** |
| | | 2002 | -0.03 | -0.04 | 0.04 | 0.32 |
| | | 2003 | -0.03 | -0.06 | 0.07 | 0.31 |
| | | 2004 | -0.07 | -0.06 | 0.06 | 0.54 |
| | | 2005 | -0.18 | -0.09 | 0.09 | 0.84 |
| | | 2006 | -0.09 | -0.05 | 0.05 | 0.77 |
| | | 2007 | -0.06 | -0.04 | 0.05 | 0.60 |
| | | 2008 | -0.04 | -0.05 | 0.05 | 0.46 |
| | | 2009 | -0.01 | -0.06 | 0.05 | 0.20 |
| | | 2010 | -0.03 | -0.07 | 0.07 | 0.29 |
| | | 2011 | -0.10 | -0.05 | 0.05 | 0.90 |
| | | 2012 | -0.07 | -0.04 | 0.04 | 0.82 |
| | | 2013 | -0.01 | -0.03 | 0.04 | 0.21 |
| | | 2014 | -0.02 | -0.03 | 0.03 | 0.47 |
| | | 2015 | -0.05 | -0.03 | 0.04 | 0.68 |
| | SU | 1994 | -0.01 | -0.05 | 0.06 | 0.22 |
| | | 1995 | -0.07 | -0.05 | 0.05 | 0.65 |
| | | 1996 | -0.04 | -0.04 | 0.07 | 0.46 |
| | | **1997** | **0.08** | **-0.05** | **0.06** | **0.02** |
| | | **1998** | **0.10** | **-0.03** | **0.04** | **<0.001** |
| | | 1999 | -0.03 | -0.02 | 0.03 | 0.64 |
| | | 2000 | -0.13 | -0.08 | 0.08 | 0.74 |
| | | 2001 | -0.11 | -0.04 | 0.05 | 0.91 |
| | | 2002 | 0.08 | -0.07 | 0.09 | 0.05 |
| | | 2003 | -0.16 | -0.13 | 0.12 | 0.61 |
| | | 2004 | -0.24 | -0.11 | 0.09 | 0.92 |
| | | 2005 | -0.04 | -0.07 | 0.07 | 0.33 |
| | | 2006 | -0.09 | -0.06 | 0.07 | 0.66 |
| | | 2007 | -0.08 | -0.06 | 0.04 | 0.67 |
| | | 2008 | -0.04 | -0.06 | 0.01 | 0.08 |
| | | 2009 | -0.05 | -0.13 | 0.05 | 0.07 |
| | | 2010 | -0.45 | -0.33 | 0.11 | 0.85 |
| | | 2011 | -0.21 | -0.10 | 0.11 | 0.83 |
| | | 2012 | -0.02 | -0.06 | 0.04 | 0.17 |
| | | 2013 | -0.06 | -0.05 | 0.05 | 0.57 |
| | | 2014 | -0.07 | -0.04 | 0.05 | 0.68 |
| | | 2015 | -0.07 | -0.08 | 0.01 | 0.24 |

| | | | | | | |
|---|---|---|---|---|---|---|
| No. pups sired | | | | | | |
| | KL | 2003 | -0.08 | -0.06 | 0.05 | 0.65 |
| | | **2004** | **0.09** | **-0.11** | **0.08** | **0.01** |
| | | 2005 | -0.35 | -0.14 | 0.17 | 0.88 |
| | | 2006 | -0.06 | -0.03 | 0.03 | 0.89 |
| | | 2007 | -0.04 | -0.04 | 0.03 | 0.54 |
| | | 2008 | 0.01 | -0.05 | 0.05 | 0.10 |
| | | 2009 | -0.14 | -0.06 | 0.05 | 0.94 |
| | | 2010 | -0.16 | -0.13 | 0.07 | 0.67 |
| | | 2011 | -0.09 | -0.06 | 0.06 | 0.69 |
| | | 2012 | -0.05 | -0.04 | 0.04 | 0.68 |
| | | 2013 | -0.05 | -0.04 | 0.03 | 0.64 |
| | | 2014 | -0.10 | -0.04 | 0.04 | 0.91 |
| | SU | 2003 | -0.07 | -0.06 | 0.04 | 0.65 |
| | | 2004 | -0.11 | -0.09 | 0.07 | 0.62 |
| | | 2005 | -0.31 | -0.14 | 0.11 | 0.94 |
| | | 2006 | -0.03 | -0.04 | 0.04 | 0.35 |
| | | 2007 | -0.17 | -0.17 | 0.06 | 0.55 |
| | | 2008 | -0.09 | -0.07 | 0.05 | 0.69 |
| | | 2009 | -0.07 | -0.10 | 0.04 | 0.24 |
| | | 2011 | -0.04 | -0.06 | 0.06 | 0.34 |
| | | **2012** | **0.10** | **-0.07** | **0.08** | **0.01** |
| | | 2013 | -0.11 | -0.06 | 0.06 | 0.75 |
| | | 2014 | -0.13 | -0.07 | 0.07 | 0.79 |
| Annual survival | | | | | | |
| | KL | 1994 | -0.05 | -0.02 | 0.02 | 0.94 |
| | | 1995 | 0.01 | -0.03 | 0.03 | 0.15 |
| | | 1996 | -0.02 | -0.02 | 0.03 | 0.52 |
| | | 1997 | -0.02 | -0.02 | 0.00 | 0.60 |
| | | 1998 | -0.01 | -0.01 | 0.02 | 0.45 |
| | | 1999 | -0.02 | -0.01 | 0.02 | 0.61 |
| | | 2000 | -0.01 | -0.01 | 0.01 | 0.53 |
| | | 2001 | -0.01 | -0.02 | 0.02 | 0.40 |
| | | 2002 | -0.05 | -0.02 | 0.02 | 0.93 |
| | | 2003 | -0.05 | -0.02 | 0.02 | 0.87 |
| | | 2004 | -0.02 | -0.03 | 0.04 | 0.35 |
| | | 2005 | 0.05 | -0.05 | 0.07 | 0.06 |
| | | **2006** | **0.02** | **-0.02** | **0.02** | **0.04** |
| | | 2007 | 0.00 | -0.02 | 0.02 | 0.22 |
| | | 2008 | 0.00 | -0.02 | 0.02 | 0.18 |
| | | 2009 | -0.05 | -0.03 | 0.03 | 0.78 |
| | | 2010 | 0.01 | -0.04 | 0.05 | 0.14 |
| | | 2011 | -0.02 | -0.02 | 0.02 | 0.52 |
| | | 2012 | -0.01 | -0.02 | 0.02 | 0.46 |
| | | 2013 | -0.02 | -0.01 | 0.02 | 0.56 |
| | | 2014 | 0.00 | -0.02 | 0.02 | 0.23 |

| | | | | | |
|---|---|---|---|---|---|
| | 2015 | 0.01 | -0.01 | 0.01 | 0.12 |
| SU | 1994 | -0.04 | -0.02 | 0.02 | 0.88 |
| | 1995 | -0.03 | -0.02 | 0.02 | 0.70 |
| | **1996** | **0.02** | **-0.02** | **0.02** | **0.03** |
| | 1997 | -0.02 | -0.02 | 0.02 | 0.47 |
| | 1998 | -0.03 | -0.02 | 0.02 | 0.70 |
| | 1999 | -0.02 | -0.01 | 0.01 | 0.72 |
| | 2000 | -0.03 | -0.01 | 0.02 | 0.78 |
| | 2001 | -0.04 | -0.02 | 0.02 | 0.78 |
| | **2002** | **0.07** | **-0.03** | **0.03** | **0.001** |
| | 2003 | -0.05 | -0.03 | 0.03 | 0.70 |
| | 2004 | -0.07 | -0.04 | 0.04 | 0.83 |
| | 2005 | -0.01 | -0.04 | 0.04 | 0.23 |
| | 2006 | 0.00 | -0.02 | 0.02 | 0.17 |
| | 2007 | -0.08 | -0.04 | 0.04 | 0.86 |
| | 2008 | -0.06 | -0.03 | 0.03 | 0.90 |
| | 2009 | -0.10 | -0.06 | 0.05 | 0.79 |
| | 2010 | 0.15 | -0.20 | 0.23 | 0.06 |
| | 2011 | -0.03 | -0.03 | 0.04 | 0.53 |
| | 2012 | -0.03 | -0.02 | 0.02 | 0.58 |
| | 2013 | 0.01 | -0.02 | 0.02 | 0.09 |
| | 2014 | 0.01 | -0.02 | 0.03 | 0.13 |
| | 2015 | -0.01 | -0.01 | 0.02 | 0.43 |

**Table S4. Results from Moran's I test demonstrating whether similarity in number of pups recruited, number of pups sired, or annual survival is related to the spatial proximity of individuals, related to STAR Methods.** Results are shown for each year/grid combination and instances of significant spatial autocorrelation are indicated in bold.

| Model | N | Fixed effect | Parameter ± SE | *z* | *P* |
|---|---|---|---|---|---|
| Survival | 475 | **Familiarity** | **0.50 ± 0.12** | **4.22** | **<0.001** |
| | | **Age** | **-0.35 ± 0.11** | **-3.17** | **0.002** |
| | | Relatedness | 0.13 ± 0.10 | 1.29 | 0.20 |
| | | Density | 0.15 ± 0.11 | 1.35 | 0.18 |
| | | Grid-SU | 0.22 ± 0.21 | 1.01 | 0.31 |
| Male ARS | 74 | **Familiarity** | **0.35 ± 0.15** | **2.35** | **0.02** |
| | | **Age** | **-0.43 ± 0.20** | **-2.12** | **0.03** |
| | | Relatedness | 0.05 ± 0.15 | 0.34 | 0.73 |
| | | Density | 0.15 ± 0.20 | 0.77 | 0.44 |
| | | Grid-SU | 0.34 ± 0.34 | 1.00 | 0.32 |
| Female ARS | 233 | **Familiarity** | **0.28 ± 0.10** | **2.75** | **0.006** |
| | | **Age** | **-0.24 ± 0.11** | **-2.18** | **0.03** |
| | | Relatedness | 0.08 ± 0.08 | 0.94 | 0.35 |
| | | **Density** | **-0.45 ± 0.17** | **-2.67** | **0.008** |
| | | **Grid-SU** | **-0.58 ± 0.21** | **-2.79** | **0.005** |

**Table S5. Fixed effects from annual survival, male annual reproductive success (ARS), and female ARS generalized linear mixed-effects models excluding years in which significant spatial autocorrelation was detected (see Table S4), related to STAR Methods.** Models shown are based on the senescent (≥ 4 years old) dataset with significant effects indicated in bold. Regression coefficients are standardized.
